# Allosteric modulation of the adenosine A_2A_ receptor by cholesterol

**DOI:** 10.1101/2021.09.13.460151

**Authors:** Shuya Kate Huang, Omar Almurad, Reizel J. Pejana, Zachary A. Morrison, Aditya Pandey, Louis-Philippe Picard, Mark Nitz, Adnan Sljoka, R. Scott Prosser

## Abstract

Cholesterol is a major component of the cell membrane and commonly regulates membrane protein function. Here, we investigate how cholesterol modulates the conformational equilibria and signaling of the adenosine A_2A_ receptor (A_2A_R) in reconstituted phospholipid bilayers. GTP hydrolysis assays show that cholesterol is a weak positive allosteric modulator of A_2A_R, as seen through enhanced basal signaling and a small decrease in agonist EC_50_. Fluorine nuclear magnetic resonance (^19^F NMR) spectroscopy suggests that this enhancement arises from an increase in the receptor’s active state populations and stronger G protein coupling. ^19^F NMR of fluorinated cholesterol analogs reveals transient and non-specific interactions with A_2A_R, indicating a lack of high-affinity binding sites or direct allosteric modulation. This is confirmed by computational analysis which suggests that cholesterol contacts confer a weak and possibly negative allosteric effect. The combined results suggest that the observed cholesterol allostery in A_2A_R is likely a result of indirect membrane effects through cholesterol-mediated changes in membrane properties, as shown by membrane fluidity measurements and high-pressure NMR.

## Introduction

In mammalian cell membranes, cholesterol accounts for ∼5-45% of the total lipid content across different cell types and subcellular components (Casares et al., 2019; Ingólfsson et al., 2017). It is a critical metabolic precursor to steroid hormones, bile salts, and vitamin D, while numerous cardiovascular and nervous system disorders are attributed to abnormalities in cholesterol metabolism (Arsenault et al., 2009; Martín et al., 2014). The rigid planar structure of cholesterol promotes ordering of bilayer lipids, thus modulating membrane fluidity and thickness. Cholesterol also drives the formation of raft-like microdomains and commonly interacts with membrane proteins as a ligand or allosteric modulator (Hulce et al., 2013).

Here, we investigate how cholesterol influences the conformational equilibria and signaling of a well-studied integral membrane protein, the adenosine A_2A_ receptor (A_2A_R), in reconstituted phospholipid/cholesterol nanodiscs. Specifically, we seek to understand if the effects on A_2A_R function are a consequence of direct allosteric interplay between cholesterol and the receptor, or if the observed effects result primarily from cholesterol-driven changes in viscoelastic properties and thickness of the lipid bilayer.

A_2A_R is a member of the rhodopsin family of G protein-coupled receptors (GPCRs). The GPCR superfamily of 7-transmembrane receptors includes well over 800 species and are targeted by over one-third of currently approved pharmaceuticals (Hauser et al., 2017). A_2A_R activates the stimulatory heterotrimeric G protein (G_s_αβγ) and is a target for the treatment of inflammation, cancer, diabetes, and Parkinson’s disease (Effendi et al., 2020; Guerrero, 2018; Ruiz et al., 2014; Yu et al., 2020; Zheng et al., 2019). Several GPCRs have been shown to interact with cholesterol, including the serotonin 5-HT_1A_ receptor, the β_2_-adrenergic receptor, the oxytocin receptor, the CCR5 and CXCR4 chemokine receptors, the CB_1_ cannabinoid receptor, and A_2A_R (Gimpl, 2016; Jafurulla et al., 2019; Kiriakidi et al., 2019). Presently, 38 out of 57 published structures of A_2A_R contain co-crystallized cholesterol (Fig. 1). In detergent preparations of A_2A_R, the soluble cholesterol analog, cholesteryl hemisuccinate (CHS), is important for receptor stability and ligand binding (O’Malley et al., 2011a, 2007). Apart from those found in crystal structures, cholesterol interaction sites within A_2A_R have also been proposed in computational studies. These include the widely conserved cholesterol consensus motif (CCM) in GPCRs, various hydrophobic patches around A_2A_R, and regions of the receptor interior (Genheden et al., 2017; Guixà-González et al., 2017; Lee et al., 2013; Lee and Lyman, 2012; Lovera et al., 2019; McGraw et al., 2019; Rouviere et al., 2017; Sejdiu and Tieleman, 2020; Song et al., 2019). The CRAC (cholesterol recognition/interaction amino acid consensus) motif, another sequence commonly found in membrane proteins that bind cholesterol, is also present in A_2A_R (Fig. 1B) (Li and Papadopoulos, 1998). Additionally, cell-based assays have shown that A_2A_R-dependent cyclic adenosine monophosphate (cAMP) production is positively correlated with membrane cholesterol (Charalambous et al., 2008; McGraw et al., 2019).

**Fig. 1.**
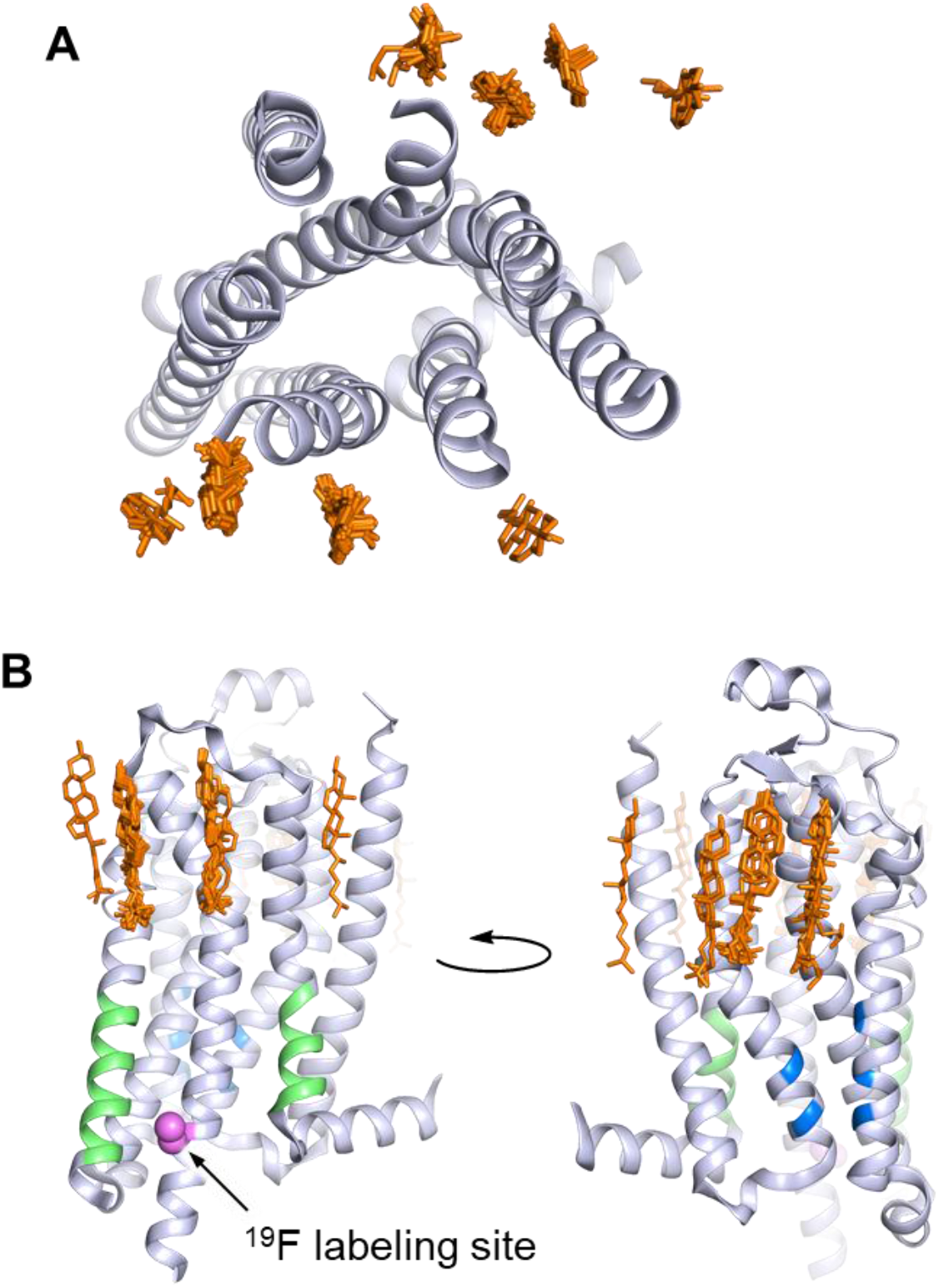
A_2A_R crystal structures reveal many cholesterol interaction sites. **(A)** Overlay of 38 currently published A_2A_R crystal structures containing co-crystallized cholesterol (extracellular view, with cholesterols shown as orange sticks). For simplicity, extracellular loops and fusion proteins are removed and only one receptor structure is shown (PDB: 4EIY). **(B)** Side views of **(A)** highlighting the CCM (blue), the CRAC motifs (green), and V229C ^19^F labeling site (violet).

Despite the prevalence of cholesterol or its analogues in many crystal structures, there is little consensus on the role that membrane cholesterol plays in A_2A_R function. While some studies found that ligand binding was unaffected by cholesterol depletion (Charalambous et al., 2008; McGraw et al., 2019), others have observed the opposite (Guixà-González et al., 2017; O’Malley et al., 2011a, 2011b). One study in particular suggested that cholesterol may laterally diffuse in the membrane and enter the receptor interior at the orthosteric site (Guixà-González et al., 2017). Additionally, whereas A_2A_R is found in both non-raft and raft-like membranes, its colocalization and modulatory effects on other cellular binding partners, including tyrosine receptor kinase B, Ca^2+^-activated K^+^ (IK1) channel, and the stimulatory G protein, have been reported to depend on cholesterol-rich microdomains (Charalambous et al., 2008; Lam et al., 2009; Mojsilovic-Petrovic et al., 2006). One possible source of discrepancy between studies is the use of different cell lines. For instance, Guixà-González *et al*. observed an increased binding by A_2A_R inverse agonist [^3^H]ZM241385 upon cholesterol depletion in C6 glioma cells. This effect was absent in a study by McGraw *et al*., who employed HEK293 cells. Cholesterol extraction or enrichment from cells exhibiting different membrane compositions and signaling patterns may trigger variable cellular response and complicates the comparison of results from different cell lines. The *in vitro* studies, on the other hand, relied on measuring ligand affinity in detergent micelles while titrating water-soluble cholesterol analogs. Although the composition of detergent preparations can be carefully controlled, the micellar environment is quite different from a lipid bilayer from the perspective of both receptor and cholesterol.

To mitigate the many complexities encountered in live cells or the inherent biases associated with detergent micelles, we employed reconstituted discoidal high density lipoprotein particles (rHDLs, also known as nanodiscs) to investigate the role of cholesterol in A_2A_R conformation and signaling. In this case, both the size and composition of these phospholipid bilayer model systems can be controlled. Through fluorine nuclear magnetic resonance spectroscopy (^19^F NMR) and *in vitro* assays, we find that cholesterol is a weak positive allosteric modulator of A_2A_R. This can be attributed to a subtle rise in population of the receptor’s active state conformers and a stronger coupling to the G protein. Interactions between A_2A_R and fluorinated cholesterol analogs appear to be short-lived and non-specific, indicating a lack of high-affinity binding sites or direct allosteric modulation. Rather, the observed allostery is likely a result of indirect membrane effects through cholesterol-mediated changes in bilayer fluidity and thickness, which can be recapitulated (without the use of cholesterol) by the application of hydrostatic pressure.

## Results

### Cholesterol is a weak positive allosteric modulator of A_2A_R

We sought to explore receptor-cholesterol allostery in a native lipid bilayer environment, free from the complexities associated with other cellular response to membrane alteration in live cells. To this end, we reconstituted A_2A_R (residues 2-317 with valine 229 mutated to cysteine for ^19^F-labelling) in nanodiscs containing a 3:2 ratio of 1-palmitoyl-2-oleoyl-*sn*-glycero-3-phosphocholine (POPC) and 1-palmitoyl-2-oleoyl-*sn*-glycero-3-phospho-(1’-*rac*-glycerol) (POPG), supplemented with different amounts of cholesterol. In our hands, cosolubilization of cholesterol with phospholipids prior to reconstitution (Midtgaard et al., 2015) resulted in polydisperse particles and low cholesterol incorporation. We therefore adapted a procedure commonly used in cells and liposomes, to deliver cholesterol via methyl-β-cyclodextrin (MβCD) to preformed nanodiscs (Zidovetzki and Levitan, 2007). This allowed us to incorporate up to ∼15 mol% cholesterol into A_2A_R-embedded nanodiscs without affecting their size distribution (Fig. S1).

To examine the effects of cholesterol on receptor-mediated G protein activation, we measured the GTPase activity of purified G proteins (G_s_α_short_β_1_γ_2_, henceforth referred to as Gαβγ) in the presence of A_2A_R-nanodiscs containing 0%, 3%, 8%, 11% and 13% cholesterol. As shown in Fig. 2A, similar agonist dose-response profiles were obtained across different cholesterol concentrations. GTP hydrolysis (cumulative over 90 min) was higher in the presence of A_2A_R relative to G protein alone and was amplified by the agonist 5’-*N*-ethylcarboxamidoadenosine (NECA) in a dose-dependent manner. Upon careful inspection, a small yet notable decrease in agonist EC_50_ values can be observed as a function of cholesterol. There is also a slight enhancement in receptor basal activity at high cholesterol concentrations (Fig. 2B-C). Thus, functionally cholesterol behaves as a positive allosteric modulator (PAM) of A_2A_R, although there is very weak cooperativity between cholesterol and agonist.

**Fig. 2.**
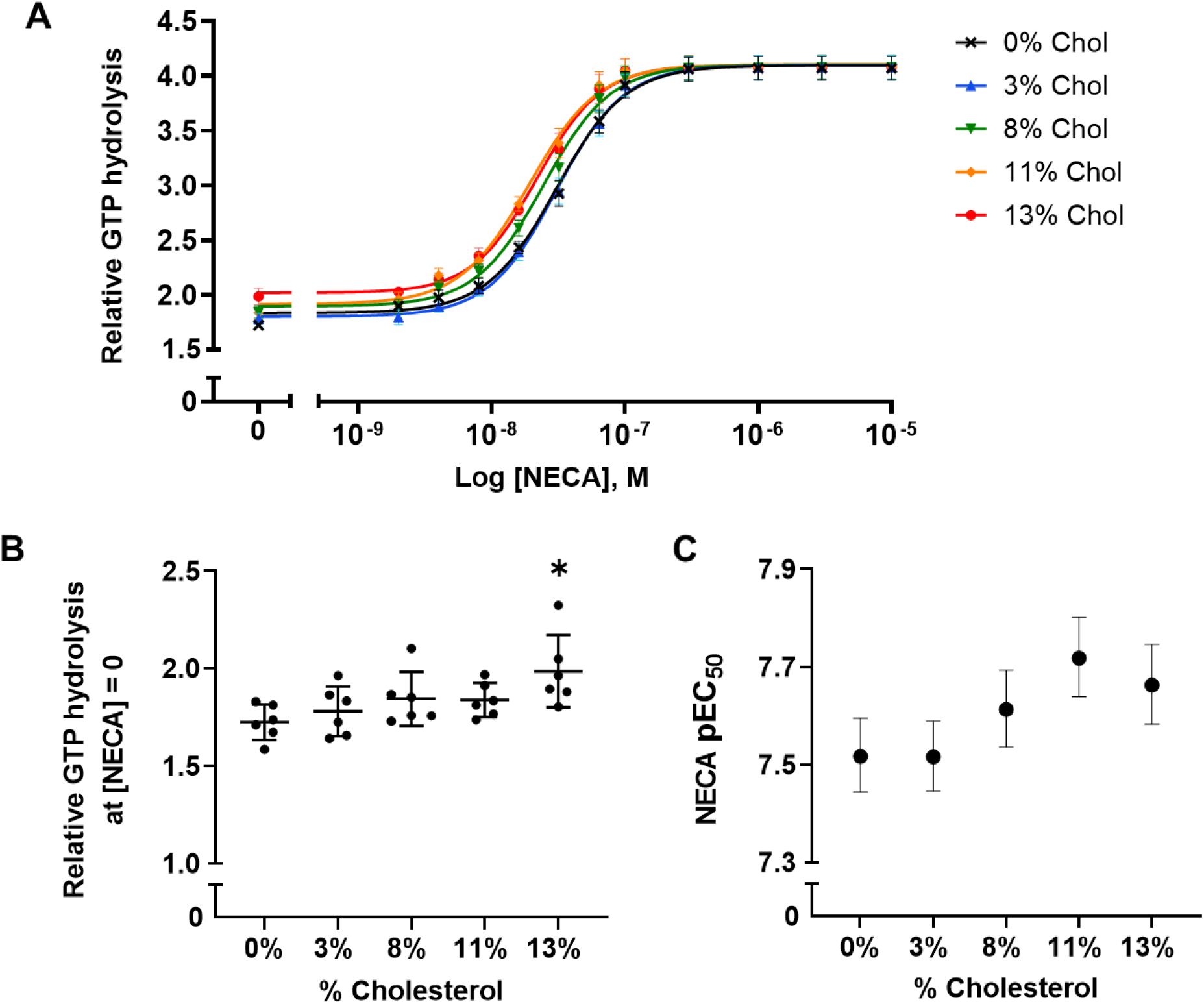
A_2A_R agonist potency and basal activity are weakly enhanced by cholesterol. **(A)** Agonist (NECA) Dose-response curves for A_2A_R-nanodiscs containing varying concentrations of cholesterol. The vertical axis represents GTP hydrolysis by purified Gαβγ (cumulative over 90 min) in the presence of A_2A_R and agonist, relative to GTP hydrolysis by Gαβγ alone in the absence of A_2A_R. Each data point represents the mean ± SEM (n = 6, technical triplicates). **(B)** Relative GTP hydrolysis in the presence of apo A_2A_R (no agonist) in nanodiscs containing varying concentrations of cholesterol. Data represents mean ± SD (n = 6, averages from each technical triplicate presented as individual points) and the asterisk represent statistical significance relative to the 0% cholesterol condition. Statistical significance was determined by one-way ANOVA followed by the Tukey’s multiple comparison test. * P ≤ 0.05. **(C)** pEC_50_ values of the NECA dose-response curves in **(A)**. Error bars represent 95% (asymmetrical profile likelihood) confidence intervals.

The weak cholesterol dependence above implies that either cholesterol does not form tight interactions with A_2A_R, or that the interactions it establishes with the receptor do not grossly overlap with the predominant allosteric pathways established by the agonist. The observed enhancement may also be a consequence of an indirect effect resulting from changes to membrane physical properties. Although the amounts of cholesterol used in this study were lower than that of a typical plasma membrane, they greatly exceed the concentrations needed to saturate potential high-affinity binding sites. Therefore, a simple allosteric mechanism involving specific binding by cholesterol is unlikely.

Using ^19^F NMR, it is possible to directly assess the effects of cholesterol on the distribution of receptor functional states. Based on the agonist dose-response curves, we expected a stabilization of activation intermediates or active states, at least in the presence of G protein. ^19^F NMR spectra of A_2A_R were recorded as a function of ligand, G protein, and cholesterol (Figs. 3A and S2). In this case, the receptor was labelled at the intracellular side of transmembrane helix 6 (TM6), in a region known to undergo large conformational changes upon activation (Fig. 3B). The resulting resonances have been assigned in our previous works and are shown as cartoons in Fig. 3C (Huang et al., 2021; Ye et al., 2016). Briefly, S_1_ and S_2_ represent two inactive state conformers differentiated by a conserved salt bridge (“ionic lock”) between TM3 and TM6. The A_3_ state is an activation intermediate stabilized by Gαβγ binding in the absence of ligands and is hence associated with the “precoupled” state. A_1_ and A_2_ represent distinct active state conformers that facilitate nucleotide exchange in the receptor G protein complex. It was found that while A_1_ is preferentially stabilized by full agonist, A_2_ is more pronounced in the presence of partial agonist.

**Fig. 3.**
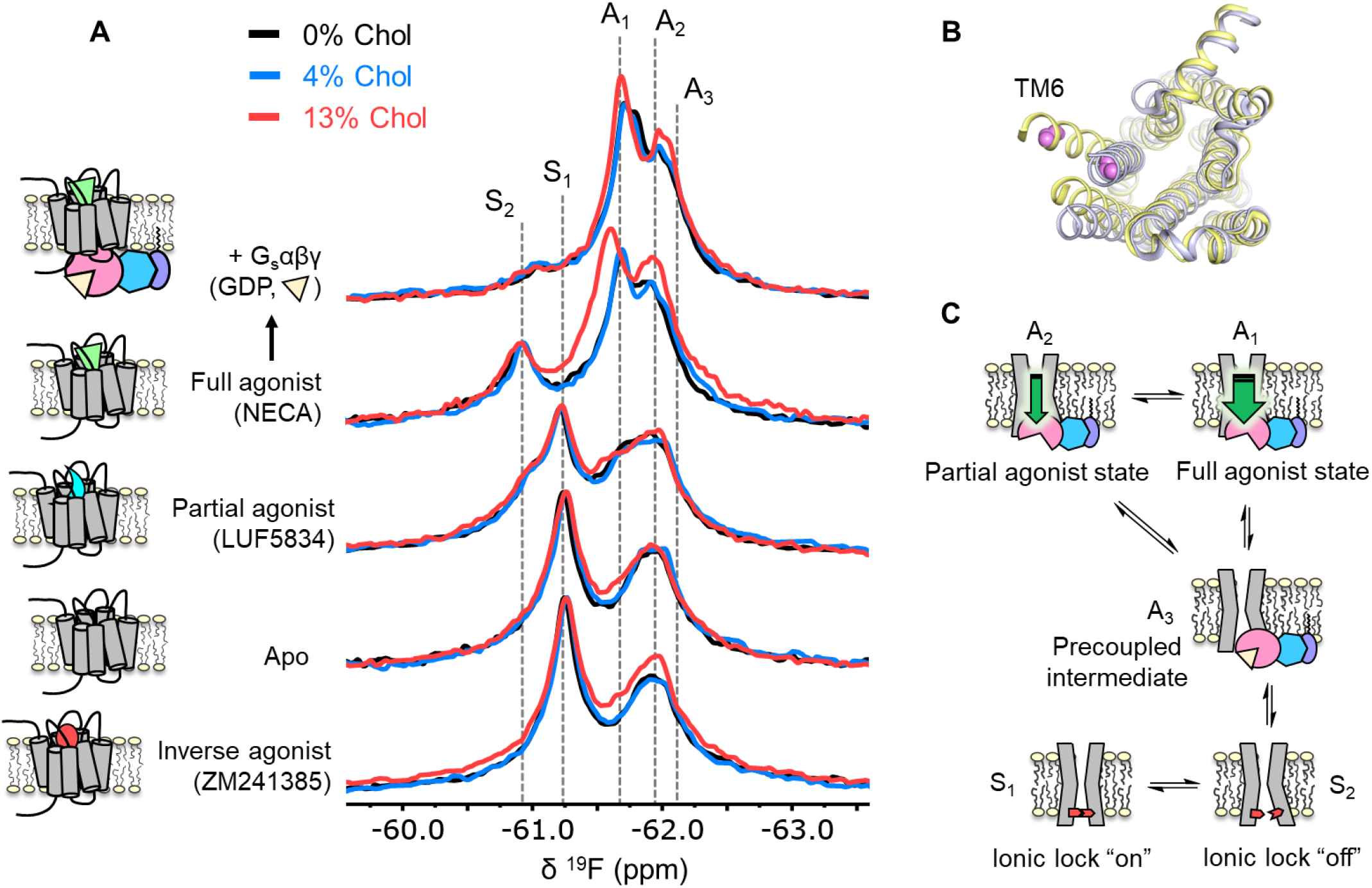
Cholesterol induces a small population increase in the active state conformers of A_2A_R. **(A)** ^19^F NMR spectra of A_2A_R in nanodiscs containing 0, 4, and 13% cholesterol, as a function of ligand and G protein. Two inactive states (S_1-2_) and three active states (A_1-3_), previously identified, are indicated by grey dashed lines. For each ligand condition, spectra from the three cholesterol concentrations are normalized via their inactive state intensity. **(B)** Intracellular view of an inactive (grey, PDB: 4EIY) and an active (yellow, PDB: 5G53) crystal structure of A_2A_R highlighting the movement of TM6 upon activation. The ^19^F-labeling site is shown in violet. **(C)** Cartoon representations of the key functional states of A_2A_R indicated in **(A)**. At the bottom are two inactive states (S_1_ and S_2_) where a conserved salt bridge is either intact or broken. A_3_ is an intermediate state that facilitates G protein recognition and precoupling. A_1_ and A_2_ are active states that drive nucleotide exchange. While A_1_ is more efficacious and preferentially stabilized by the full agonist, A_2_ is less efficacious and reinforced by a partial agonist.

Inspection of the overlaid spectra in Fig. 3A reveals nearly identical distributions of conformational states between A_2A_R in the presence of 0% and 4% membrane cholesterol. At 13% cholesterol, subtle changes can be observed in the inverse agonist-saturated, apo, full agonist-saturated, and full agonist + Gαβγ spectra. In particular, we observed a small population shift toward the active states, A_1_ and A_2_. The results imply that cholesterol is a positive allosteric modulator of A_2A_R and acts in part through stabilization of the active state ensemble. Interestingly, 13% cholesterol resulted in line-broadening and a 0.09 ppm downfield shift of the A_1_ state in the full agonist-saturated spectrum, changes which are not observed in the full agonist + Gαβγ condition. The changes seen with the A_1_ resonance in the presence of 13% cholesterol and NECA alone could be a consequence of motional averaging and/or a perturbation of the average orientation of TM6. Clearly, cholesterol exerts differential effects on distinct conformers in the ensemble. The outcome of the NMR experiments is consistent with the GTPase activity assay. Nearly complete overlap between the 0% and the 4% spectral series suggest that the principal allosteric mechanism is unlikely related to high-affinity binding. While the subtle changes observed at 13% cholesterol are evidence for positive allosteric modulation, the effects are much smaller than those of any orthosteric ligands or other known allosteric modulators of A_2A_R (Gao et al., 2020; Ye et al., 2018).

To understand if the weakly activating role of cholesterol arises because of enhanced efficiency in nucleotide exchange or pre-association with G protein (precoupling), we carried out ^19^F NMR experiments on apo-A_2A_R in the presence of Gαβγ without agonist. As mentioned above, this condition produces the precoupled receptor-G protein complex and greatly stabilizes the A_3_ state (Huang et al., 2021). This is recapitulated in Fig. 4 for all three cholesterol concentrations, where a shift in the equilibrium populations toward the active conformers, particularly A_3_ and A_2_, is observed upon the addition of Gαβγ. Importantly, an increase in cholesterol further enhanced the A_3_ state in addition to a decrease in the peak width. The magnitudes of these changes are small, consistent with results shown in Figs. 2-3. The results suggest that membrane cholesterol may help to stabilize the precoupled complex of A_2A_R and G protein, and possibly modulates the amplitudes of motion about the precoupled state. This in turn may favor further conformational exchange to A_1_ or A_2_. Taken together, ^19^F NMR showed that mechanistically, the PAM effect of cholesterol in A_2A_R can be attributed to an increase in the population of active state conformers as well as a more robust coupling to the G protein.

**Fig. 4.**
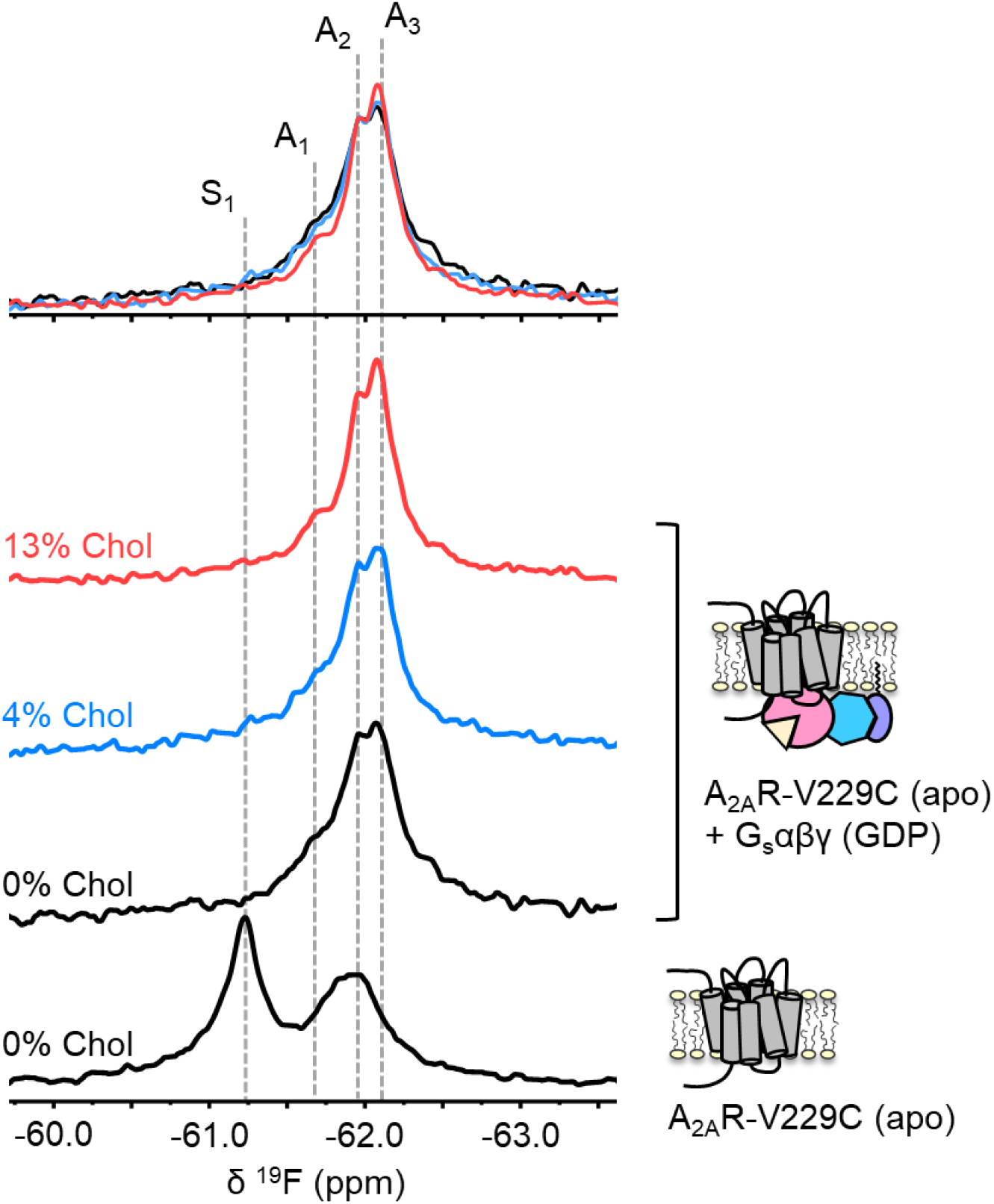
The precoupled complex of A_2A_R-G_s_αβγ is stabilized by cholesterol. ^19^F NMR spectra of apo A_2A_R in the presence of 0, 4, and 13% cholesterol, and as a function of Gαβγ. The key functional states are indicated by grey dashed lines and the three spectra in the presence of G protein are normalized via the A_2_ state.

### Allosteric network analysis reveals small negative allosteric modulation by cholesterol

Given the above observations, we employed rigidity-transmission allostery (RTA) analysis (Sljoka, 2021) to survey allosteric activation pathway perturbation by cholesterol within the ternary complex. The RTA algorithm is a computational tool based on mathematical rigidity theory and has been used to identify allosteric networks within proteins (Jacobs et al., 2001; Sljoka, 2021; Whiteley, 2005). It predicts how changes in the conformational rigidity or flexibility of one region in the protein are transmitted to distal sites by quantifying the resulting differences in the degrees of freedom within the system. Similarly, ligand-induced perturbations can be examined by rigidifying the ligand itself or its binding pocket. Using a model of an agonist- and GDP-bound A_2A_R-G_s_αβγ complex, equilibrated in a 1 µs simulation in POPC bilayer with 20% cholesterol, our previous work revealed that rigidification of the agonist NECA results in changes in the degrees of freedom which can be transmitted from the orthosteric pocket to the Gα nucleotide binding site (Huang et al., 2021). This allosteric network encompasses large portions of the receptor, the N- and C-terminal helices of Gα, parts of the Gα Ras domain, three out of seven beta propellers within Gβ, and a section of the Gβ N-terminal helix that forms coiled-coil interactions with Gγ.

Using the above model, we repeated the RTA analysis to examine whether this previously identified allosteric network is sensitive to the presence of cholesterol. Seven cholesterol molecules were found in the vicinity (within 6 Å) of A_2A_R and were removed prior to rigidification of the agonist. The resulting change in degrees of freedom is mapped in Fig. 5 for each residue within the ternary complex. Removal of cholesterol gave rise to an allosteric pathway which is very similar to that in the presence of cholesterol, although with altered intensities for some regions. Higher allosteric transmission is observed for the CWxP motif of TM6, in particular the tryptophan toggle switch W246^6.48^, in the absence of cholesterol. On the other hand, stronger allosteric transmission is observed for the NPxxY motif of TM7 in the presence of cholesterol. Interestingly, the removal of cholesterol resulted in a slight overall enhancement in allosteric transmission to the G protein. This includes the Gα N- and C-terminal helices which interact with the receptor, as well as Gβ which has been found to play a role in conferring ligand efficacy (Huang et al., 2021). The above observations suggest that the overall presence of cholesterol, while not drastically perturbing, reduces signal transmission across the ternary complex. This is inconsistent with our experimental observations, however, which indicate that cholesterol is a weak PAM.

**Fig. 5.**
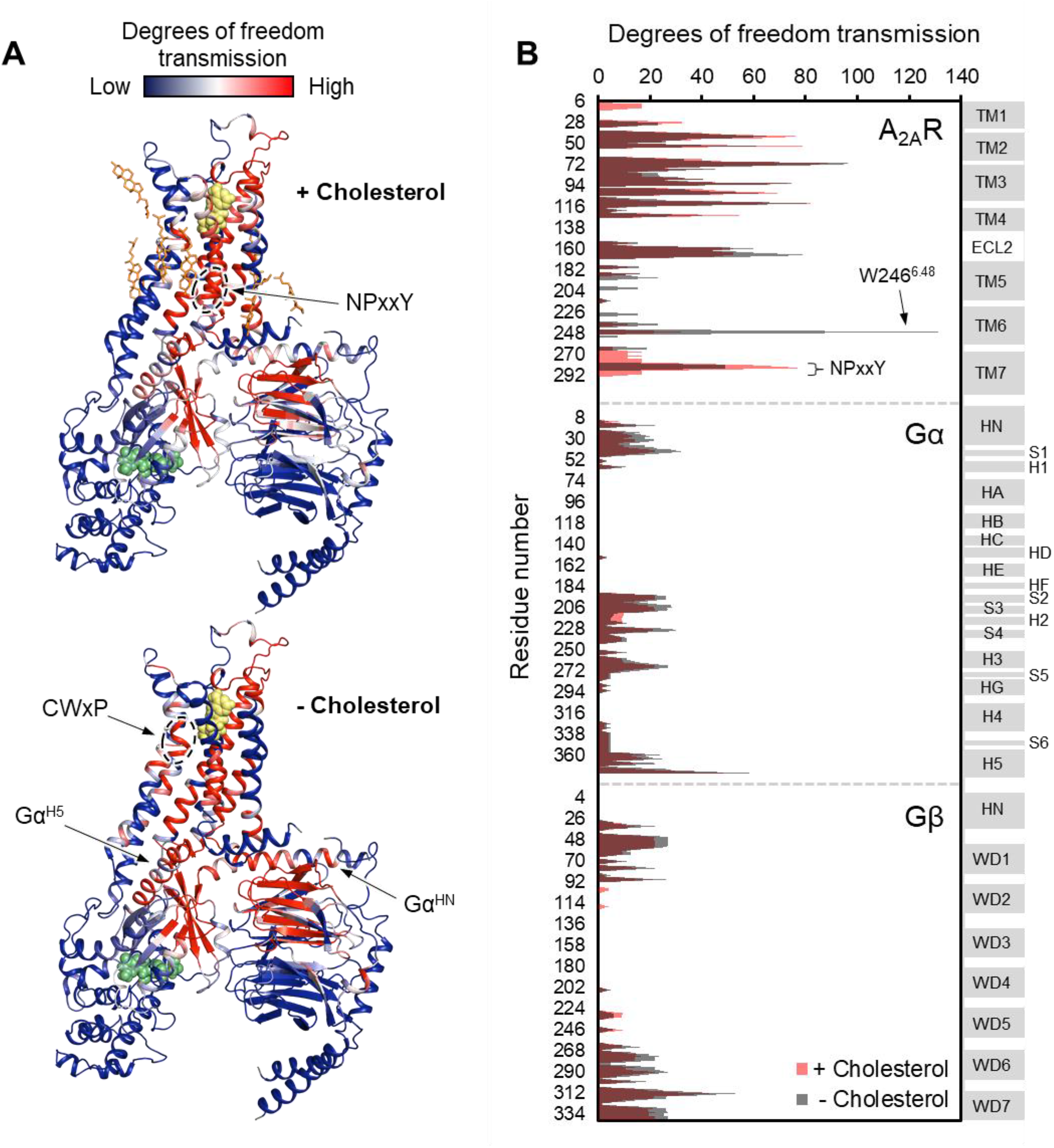
Removal of cholesterol leads to a small overall enhancement in allosteric transmission from the agonist binding site. **(A)** Allosteric networks within the A_2A_R-Gαβγ complex in the presence and absence of cholesterol, revealed through RTA analysis via rigidification of the agonist NECA (yellow spheres). The intensity of allosteric transmission is measured by the resulting regiospecific changes in degrees of freedom and is mapped in color (red/blue gradient bar). Cholesterol molecules are shown as orange sticks while green spheres represent GDP. **(B)** The intensity of allosteric transmission is plotted for each residue in A_2A_R, Gα, and Gβ. Secondary structural elements are indicated on the right. Grey blocks denote α-helices and β-strands, while white gaps represent loops. For the Gα subunit, the common Gα numbering system is used (Flock et al., 2015).

### A_2A_R-cholesterol interactions are short-lived and non-specific

To further evaluate the nature of cholesterol-A_2A_R interactions, we carried out ^19^F NMR experiments of fluorinated cholesterol analogs, delivered into either empty or A_2A_R-embedded nanodiscs via MβCD. Two different molecules were tested (Fig. 6). 3β-fluoro-cholest-5-ene (3β-F-chol) was synthesized in house and features a fluorine atom in place of the cholesterol hydroxyl headgroup. The fluoro group is a relatively benign substitute for the hydroxyl due to its similar size and electronegativity. It also retains some ability to accept hydrogen bonds (Hoffmann and Rychlewski, 2002). Another cholesterol analog, referred to as F7-chol, was purchased commercially and had the tail isopropyl group replaced by CF(CF_3_)_2_. Incorporation of these analogs did not affect the response of A_2A_R toward ligands nor its ability to activate the G protein (Fig. S3).

**Fig. 6.**
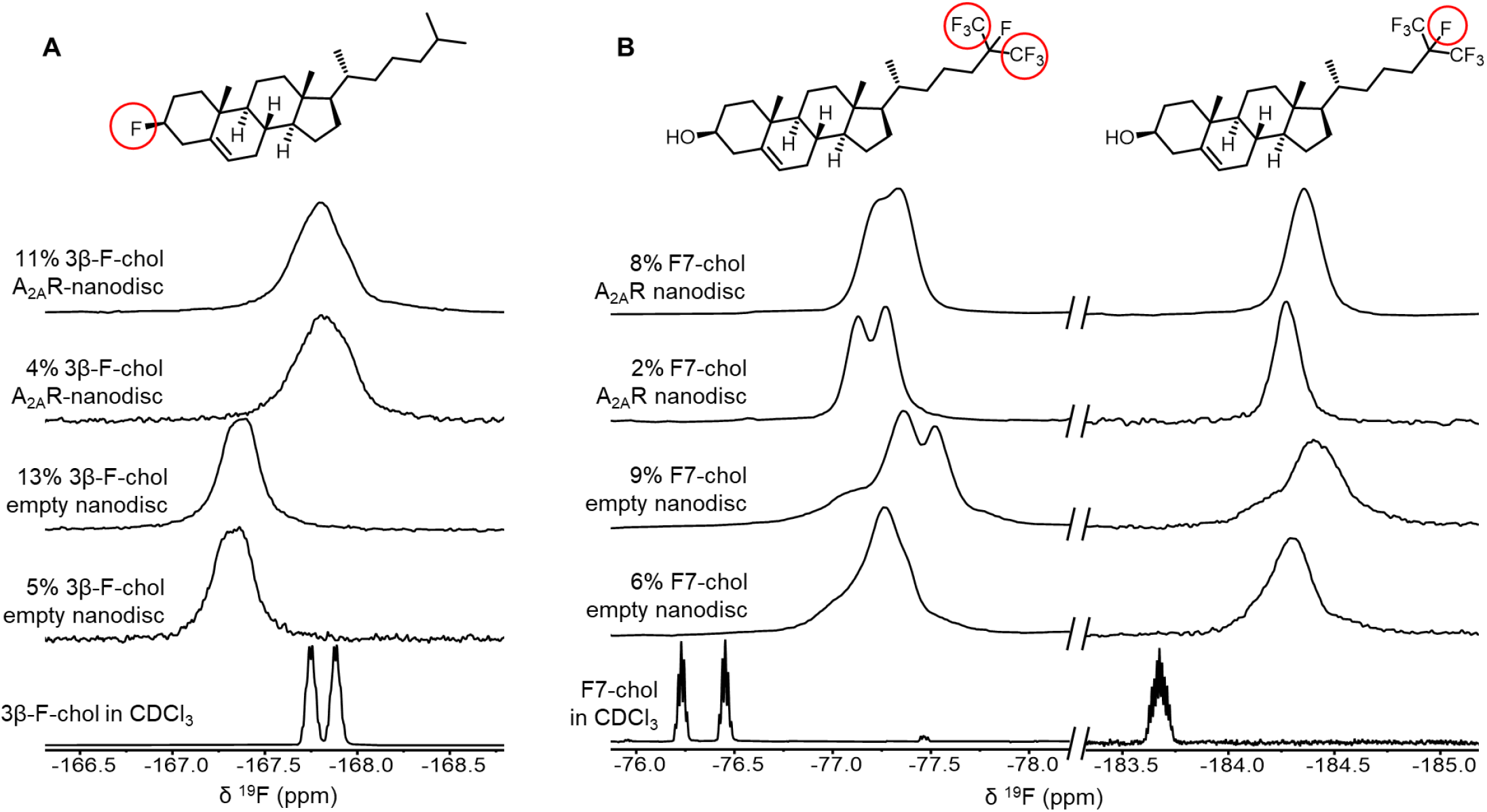
^19^F-cholesterol analogs interact with A_2A_R but show no evidence of a long-lived bound state. Non-decoupled ^19^F NMR spectra of 3β-F-chol **(A)** and F7-chol **(B)** in chloroform, empty nanodisc, and A_2A_R (apo)-embedded nanodisc. The fluorine groups contributing to each of the resonances are circled and shown above the corresponding peak.

The NMR resonances were significantly broadened for both cholesterol analogs upon incorporation into the membrane (Fig. 6). This is expected for lipid molecules situated in a slow-tumbling nanodisc. In the case of F7-chol, the peak shapes are further complicated by resonance overlap of the two CF_3_ groups, which are inequivalent and exhibit complicated multiplicity patterns. Comparison between the spectra of 3β-F-chol in empty nanodiscs and A_2A_R-embedded nanodiscs shows a clear environmental difference in the presence of receptor (Fig. 6A). The resonance is ∼0.5 ppm upfield shifted and broader relative to empty nanodiscs. However, the lack of chemical shift difference between 4% and 11% 3β-F-chol suggests that the above changes are predominantly a result of altered environment (*i*.*e*. availability of hydrophobic proteinaceous surfaces) rather than a shift toward receptor-bound states at specific binding pockets.

The spectra of F7-chol are harder to interpret. Due to the two overlapped CF_3_ resonances, a small change in chemical shift for either one could bring about dramatic variation in the overall peak shape (Fig. 6B). As such, we cannot be confident about whether the observed changes in the CF_3_ peaks in response to A_2A_R are a consequence of specific binding. However, it is clear from the two empty nanodisc spectra (containing either 6% or 9% F7-chol) that the membrane environment is altered with increasing F7-chol. Overall, the NMR data from the two ^19^F-cholesterol analogs show environmental differences between empty and A_2A_R-embedded nanodiscs as well as between different cholesterol concentrations. However, there is no direct evidence of a long-lived receptor-bound state.

As a classical PAM, cholesterol would be expected to exhibit stronger binding to A_2A_R in the presence of an agonist or G protein, versus an inverse agonist. The opposite would hold for a classical negative allosteric modulator (NAM). Yet, there was no apparent difference in chemical shift sensitivity toward agonist or inverse agonist for either ^19^F cholesterol analogs (Fig. S4). The NMR spectra of 3β-F-chol in A_2A_R-embedded nanodiscs are nearly identical upon the addition of inverse agonist, full agonist, and mini-G, a G protein mimetic that has been shown to stabilize the A_1_ active state (Carpenter et al., 2016; Huang et al., 2021). Small chemical shift changes were observed for F7-chol between the apo receptor and the ligand/mini-G-bound conditions. However, the direction of shift is the same between full agonist and inverse agonist. Thus, cholesterol interactions are independent from the identity of the orthosteric ligand bound to A_2A_R, despite being observed as a functional PAM *in vitro* (Figs. 2-3) and predicted as a NAM *in silico* (Fig. 5). This leads us to consider that the observed positive allosteric effects of cholesterol are predominantly indirect and relayed through the physical changes to the membrane bilayer.

### Cholesterol allostery in A_2A_R may be a result of indirect membrane effects

The effects of cholesterol on the physical properties of lipid bilayers have been well documented. The planar structure of cholesterol promotes orientational order in the liquid disordered phase of the bilayer, leading to reduced lateral diffusion and increased hydrophobic thickness (Fig. 7A) (Crane and Tamm, 2004; De Meyer and Smit, 2009; Filippov et al., 2003; Hung et al., 2007). For instance, as much as 20% increase in thickness can be expected for a POPC bilayer when the cholesterol concentration is varied from 0 to 30% (Mouritsen and Bagatolli, 2016; Tharad et al., 2018).

**Fig. 7.**
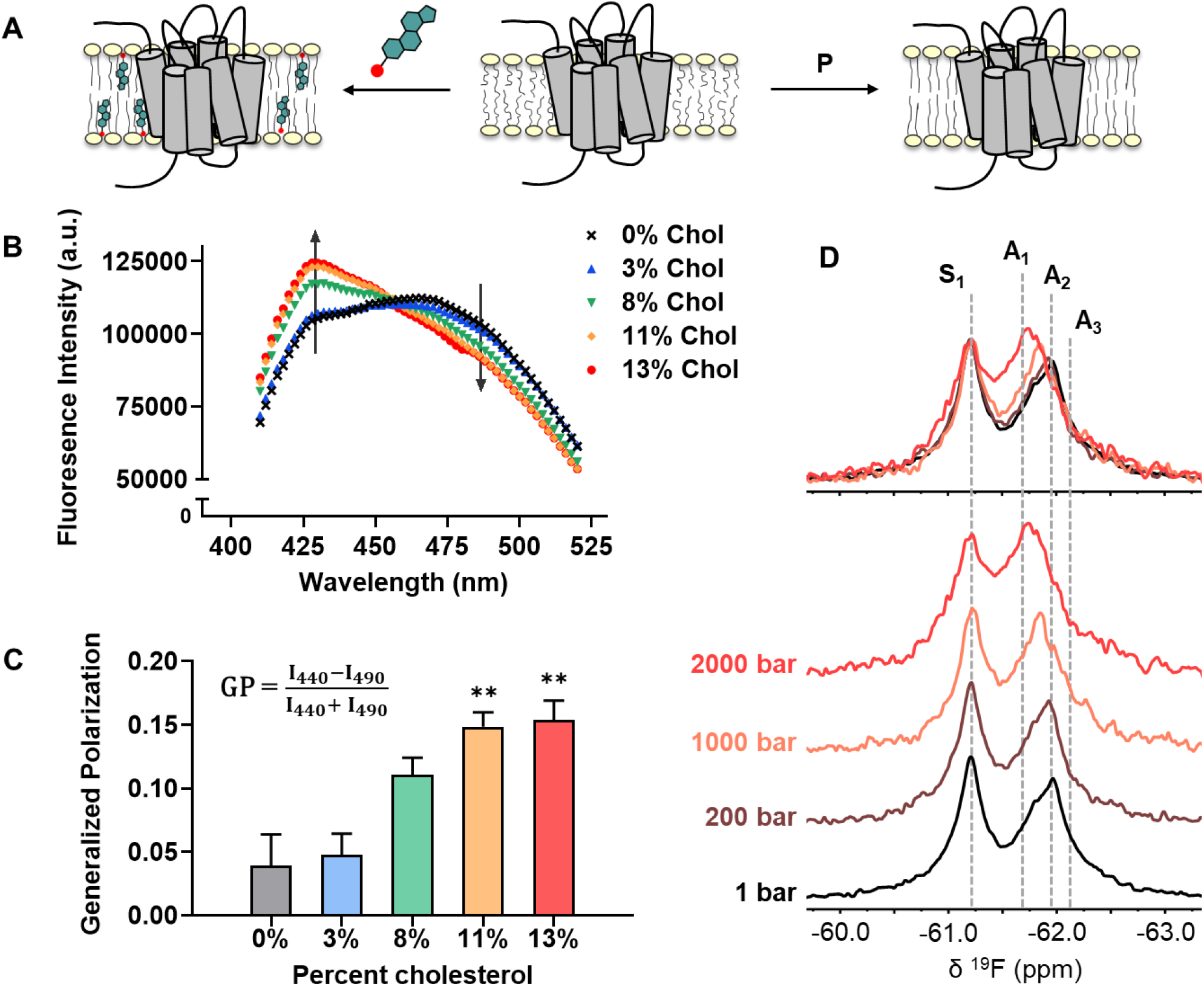
Cholesterol allostery in A_2A_R is likely a result of indirect membrane effects. **(A)** The lipid bilayer can be rigidified and thickened upon addition of cholesterol or an increase in lateral pressure. **(B)** Averaged fluorescence spectra (n = 4) of Laurdan in A_2A_R-embedded nanodiscs containing varying concentrations of cholesterol. **(C)** The emission intensity of Laurdan at 440 nm and 490 nm were used to calculate the generalized polarization values. Data represent mean ± SEM (n = 4, technical triplicates). Astrisks represent statistical significance over both the 0% and the 3% conditions. Statistical significance was determined via one-way ANOVA followed by Tukey’s multiple comparisons test. ** P ⩽ 0.01. **(D)** ^19^F NMR spectra of A_2A_R, in the absence of ligand or cholesterol, acquired at 1, 200, 1000, and 2000 bar pressures. The key functional states are inducated by grey dashed lines.

We employed the lipophilic fluorescent probe Laurdan to monitor the membrane orientational order of A_2A_R-embedded nanodiscs as a function of cholesterol. The generalized polarization 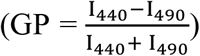 of Laurdan fluorescence is a consequence of solvent contact. Fluid (disordered) membranes exhibit smaller GP values, a consequence of dipole relaxation between Laurdan and nearby water molecules which causes a red shift of the emission wavelength. In a phospholipid bilayer, greater water accessibility at the hydrophobic-hydrophilic interface is typically a signature of enhanced reorientational dynamics within the lipid milieu (Yu et al., 1996). As shown in Fig. 7B-C, we observed a consistent shift of the Laurdan fluorescence spectra which gave rise to higher GP values at elevated cholesterol concentrations. This indicates that cholesterol incorporation enhances lipid orientational order in A_2A_R-embedded nanodisc bilayers.

An enhanced orientational order arises from a higher fraction of trans conformers in the lipid chains and consequently, an increased hydrophobic thickness. In the case where A_2A_R adopts an ensemble of states, the equilibrium is expected to shift toward those states which are more compatible with an increased hydrophobic thickness (Andersen and Koeppe, 2007). Bilayer thickness can be readily modulated by changing the composition of lipids or acyl chain lengths. Alternatively, the application of hydrostatic pressure can be used in an NMR experiment to affect changes in bilayer properties while avoiding potential complications associated with specific lipid-receptor interactions. For a typical liquid disordered phosphatidylcholine bilayer, pressure-induced compression is far more significant in the lateral than in the transverse direction (Stamatoff et al., 1978). An elevation in pressure at constant temperature promotes ordering of the fatty acyl chains. This leads to an increase in lipid packing density and hydrophobic thickness, and a reduction of lateral diffusion (Ding et al., 2017). Thus, hydrostatic pressure provides an effective way to mimic the effects of cholesterol on a lipid bilayer (Fig. 7A).

We recorded the ^19^F NMR spectra of the apo receptor in nanodiscs without cholesterol, at a pressure of 1, 200, 1000, and 2000 bar. Like cholesterol, the rise in pressure resulted in a bias toward the active ensemble, particularly the A_1_ (full agonist) state (Fig. 7D). Interestingly, the magnitude of change is non-linear and considerably larger at 2000 bar in comparison to 1000 and 200 bar. This is consistent with the expected change in membrane thickness as a function of pressure. For a pure POPC bilayer at 20 °C, the increase in hydrophobic thickness is small (up to ∼2 Å) and roughly linear below 1200 bar. Above this pressure, the bilayer transitions to a solid ordered phase which results in a rapid increase of membrane thickness on the order of 10 Å (Rappolt et al., 2003). In comparison, the thickness increase as a result of 10-15% cholesterol is on the order of 1-3 Å (Hung et al., 2007).

The NMR results are similar to that of previous pressure studies of the β_1_AR and β_2_AR in detergent micelles (Abiko et al., 2019; Lerch et al., 2020). In both cases a shift toward the active state was observed in response to pressure, which was correlated with a reduction in void volume of the active receptor relative to the inactive form. Here, ^19^F NMR allowed a more detailed delineation of the conformational landscape of A_2A_R. Unlike agonist- or G protein-induced activation, where the inactive ensemble is significantly diminished and all three active state conformers are promoted, the redistribution of states brought about by pressure saw a smaller decrease of the inactive ensemble and a specific shift in equilibrium toward the A_1_ state (Fig. 7D). The effects from pressure directly exerted on the receptor cannot be easily separated from indirect effects that are manifested through changes in the lipid bilayer. However, more influence from the membrane is expected since the molecular assembly of lipid bilayers is much more sensitive to pressure relative to the conformation of proteins (Michiko and Rikimaru, 1999). The lipid bilayer, relative to detergent micelles, was shown to protect integral membrane proteins from pressure-induced denaturation (Kangur et al., 2008). Overall, our pressure-resolved NMR data suggest that A_2A_R can be regulated indirectly through changes in the lipid bilayer. While the mechanism may be complex and the effects are subtle, receptor activation appears to be favored in an environment with higher packing density, acyl chain order, and hydrophobic thickness.

## Discussion

A_2A_R has been intensely studied by both X-ray crystallography and more recently by electron cryomicroscopy (cryo-EM). In many cases, cholesterol or CHS have proven useful in stabilizing the receptor and obtaining high resolution structures. Earlier *in vitro* and cell-based studies, along with the clear delineation of cholesterol in many crystal structures of A_2A_R suggest that the molecule may play a direct allosteric role in modulating receptor function. A body of computational work has since showcased cholesterol hot spots across the receptor and some of these studies proposed state-dependent interactions (Lovera et al., 2019; McGraw et al., 2019). Nevertheless, there is no literature consensus on the allosteric role of cholesterol on this prototypical GPCR.

In this study, we set out to investigate both the magnitude and origin of the allosteric interplay between cholesterol and A_2A_R in phospholipid bilayers. Functional and spectroscopic studies in nanodiscs identify cholesterol as a weak PAM. Specifically, GTP hydrolysis assays found a marginal increase in basal activity with increasing cholesterol, in addition to a very weak enhancement in the agonist potency. ^19^F NMR experiments revealed little or no difference in the receptor spectra upon addition of 4% cholesterol. A very modest shift in equilibrium toward the active states (A_1_ and A_2_) was observed at 13% cholesterol, indicating weak positive allostery. A distinct enhancement of A_3_ is also found at 13% cholesterol for apo receptor bound to G protein, implying that cholesterol either directly or indirectly stabilizes the precoupled A_2A_R-Gαβγ complex. However, further computational analysis predicted that the presence of cholesterol reduces allosteric transmission within the ternary complex, suggesting negative allostery by cholesterol.

In seeking an explanation for the observed positive allosteric effects of cholesterol on A_2A_R, we first considered the possibility of a bound state through ^19^F NMR experiments using fluorinated cholesterol analogs. However, no apparent binding isotherm could be observed, implying a weak or transient interaction between cholesterol and A_2A_R in phospholipid nanodiscs. There is also no correlation between the chemical shifts of the cholesterol analogs and orthosteric ligand efficacy, suggesting that the origin of the observed positive allostery is through the indirect effects of cholesterol on the membrane itself. Laurdan fluorescence experiments confirmed that lipid orientational order is indeed increased by cholesterol. Furthermore, an increased hydrostatic pressure predicted to yield comparable changes in membrane fluidity and thickness as those seen with cholesterol gave rise to a shift in receptor equilibrium toward the fully active state. There may indeed be subtle NAM effects from direct interaction with cholesterol, as suggested by our computational analysis, which at the same time are overcome by stronger indirect effects through the membrane.

While we cannot rule out the possibility of a cumulative influence from multiple fast-exchanging, weakly binding interactions, results from the current study strongly suggest that changes in membrane physical properties are the primary, albeit indirect means by which cholesterol regulates A_2A_R. It is possible that this is also the mechanism through which CHS enhances the ligand binding activity of A_2A_R in detergent micelles (*i*.*e*. by modulating the micellar structure to a more bilayer-like morphology). In support of this idea, an evaluation of A_2A_R reconstituted in various mixed micelle systems revealed a correlation between receptor activity to those detergent/CHS compositions that gave rise to a micellar hydrophobic thickness that closely matches that of native mammalian bilayers (O’Malley et al., 2011b).

One limitation of the current study is the range of cholesterol concentrations being probed, which is below the physiological norm. In cell-based experiments, total cholesterol depletion is not possible without adversely affecting cellular integrity. In many cases, the amount of cholesterol left in the membrane was not quantified and the focus was instead on the disruption of raft-like domains. Our nanodisc samples contained 0-13% cholesterol, which is below the concentration regime for raft formation (Barrett et al., 2013; Crane and Tamm, 2004). These two strategies (extraction from cholesterol-rich membranes and delivery into cholesterol-free membranes) explore largely different processes; the former involves the disruption of rafts while the latter allows studies of the interaction of cholesterol monomers with the receptor.

Our data suggests that such interactions, if present for A_2A_R, are non-specific and short-lived. This may explain why structural and computational work has yet to converge upon a single cholesterol binding site. Like lipids, the observation of cholesterol in crystal structures may simply be a consequence of having cholesterol as a part of the crystallization matrix. In fact, many A_2A_R structures which do not contain co-crystallized cholesterol (all the active state structures and some inactive state structures) had the molecule present in large quantities during crystallization. In one example, complexes of A_2A_R bound to an engineered mini-G protein were crystallized in octylthioglucoside micelles either in the presence or absence of CHS. No discernible difference was found between crystals that grew with or without CHS and the structure was solved using data collected from two crystals, one from each condition (Carpenter et al., 2016). Similarly, the numerous cholesterol “hot spots” predicted through computational approaches may not necessarily indicate functional specificity, but rather geometric compatibility between certain hydrophobic patches or grooves surrounding the receptor and the cholesterol backbone. This is reflected in the fact that nearly all seven transmembrane helices and grooves between helices in A_2A_R have been predicted in various studies to bind cholesterol (Genheden et al., 2017; Guixà-González et al., 2017; Lee and Lyman, 2012; Lovera et al., 2019; McGraw et al., 2019; Rouviere et al., 2017; Sejdiu and Tieleman, 2020; Song et al., 2019). Furthermore, the presence of CCM or CRAC motifs has recently been shown to not be predictive of cholesterol binding in GPCRs (Taghon et al., 2021).

The current work shows that A_2A_R does not require cholesterol to function in an *in vitro* bilayer setting. However, many experiments have highlighted the role of cholesterol-rich domains for A_2A_R to function in a cellular context. As alluded to above, a major shortcoming of our nanodisc system is the upper limit of cholesterol that can be delivered. This prevented us from evaluating the system at higher, more physiological cholesterol concentrations and probing the effects from protein partitioning between liquid ordered and liquid disordered phases (Gutierrez et al., 2019). It is unclear whether A_2A_R alone prefers certain regions on the plasma membrane. Nevertheless, both the stimulatory G protein and many isoforms of adenylyl cyclase were shown to partition into raft-like domains (Kamata et al., 2008; Oh and Schnitzer, 2001; Ostrom et al., 2001; Ostrom and Insel, 2004). Spatial co-localization of the receptor with other cellular binding partners in these membrane regions may therefore be required to form and maintain signaling complexes.

## Supplementary Figures

**Fig. S1.**
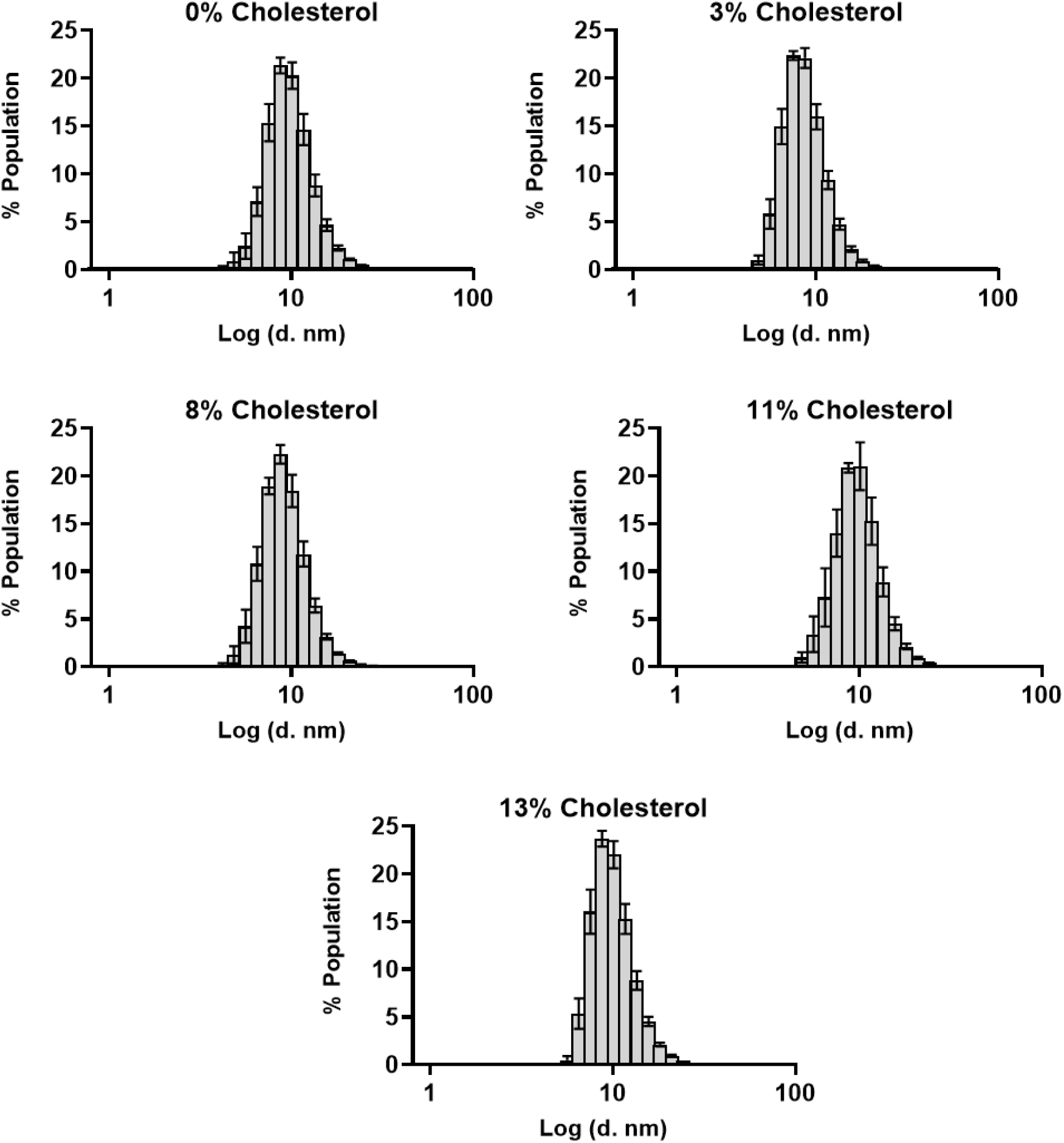
The size distribution of nanodiscs are minimally affected by cholesterol incorporation. Hydrodynamic diameters of A_2A_R-nanodiscs containing varying levels of cholesterol, measured through dynamic light scattering.

**Fig. S2.**
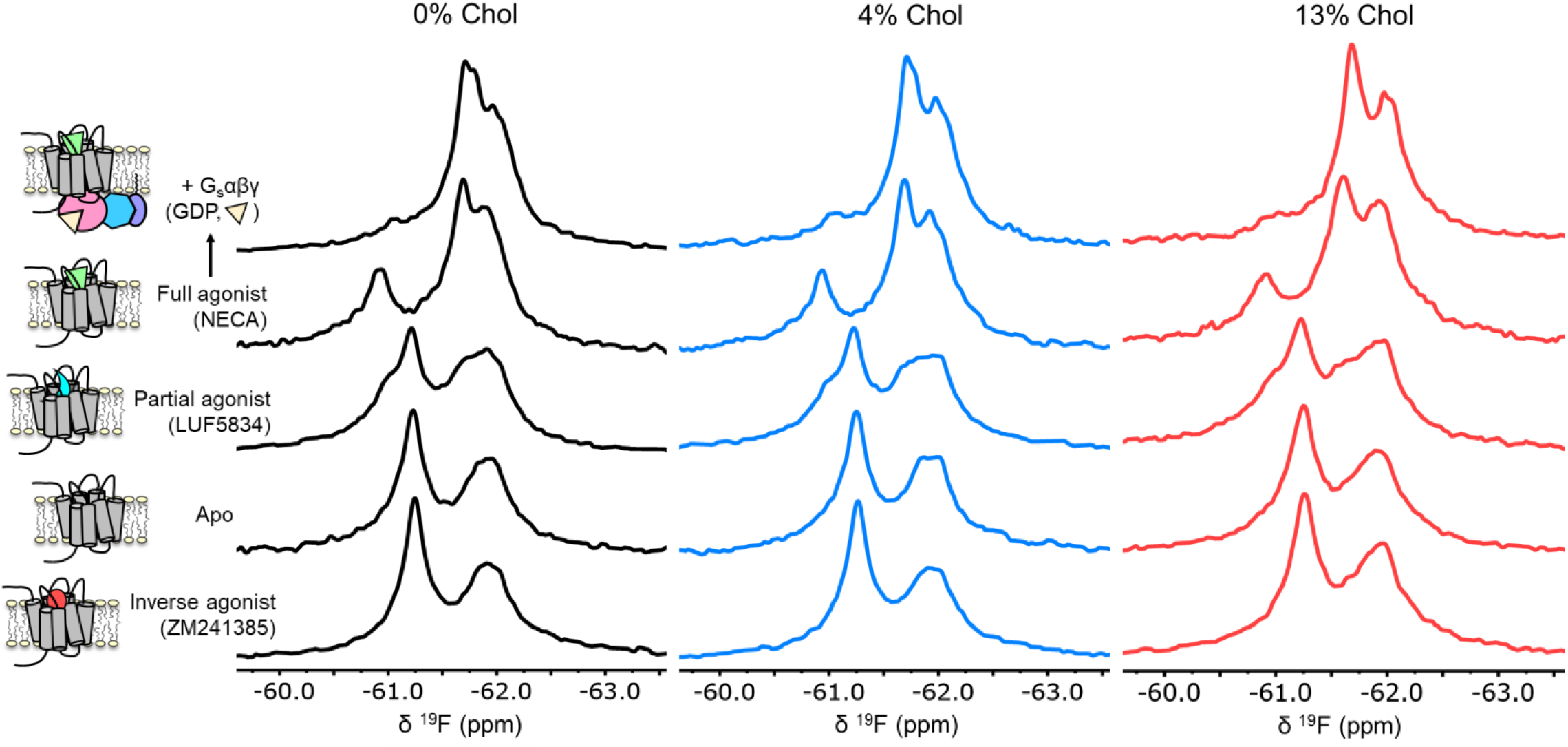
A_2A_R exhibits similar activation signatures in the absence and presence of cholesterol. Non-overlapped (relative to **Fig. 3**) ^19^F NMR spectra of A_2A_R-V229C in nanodiscs containing 0, 4, and 13% cholesterol.

**Fig. S3.**
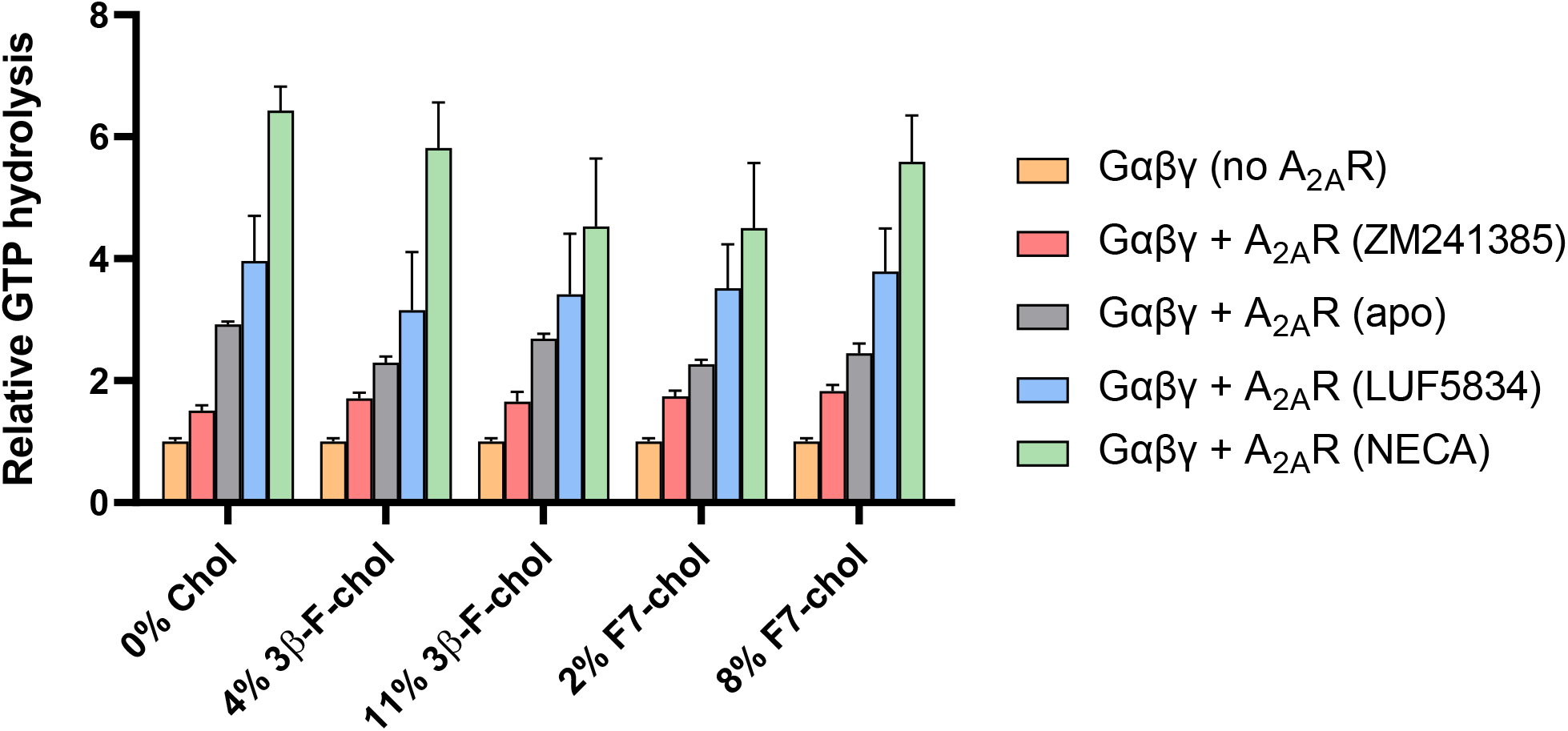
Incorporation of ^19^F-cholesterol analogs into nanodiscs did not affect A_2A_R ligand sensitivity and G protein activation. Cumulative hydrolysis of GTP by Gαβγ in the presence of A2AR-nanodiscs with and without ^19^F-cholesterol analog, relative to GTP hydrolysis by Gαβγ alone in the absence of A2AR. To assess ligand sensitivity, samples were saturated with either inverse agonist (ZM241385), no ligand, partial agonist (LUF5834), or full agonist (NECA). Data represent mean ± SEM (n = 3, technical triplicates).

**Fig. S4.**
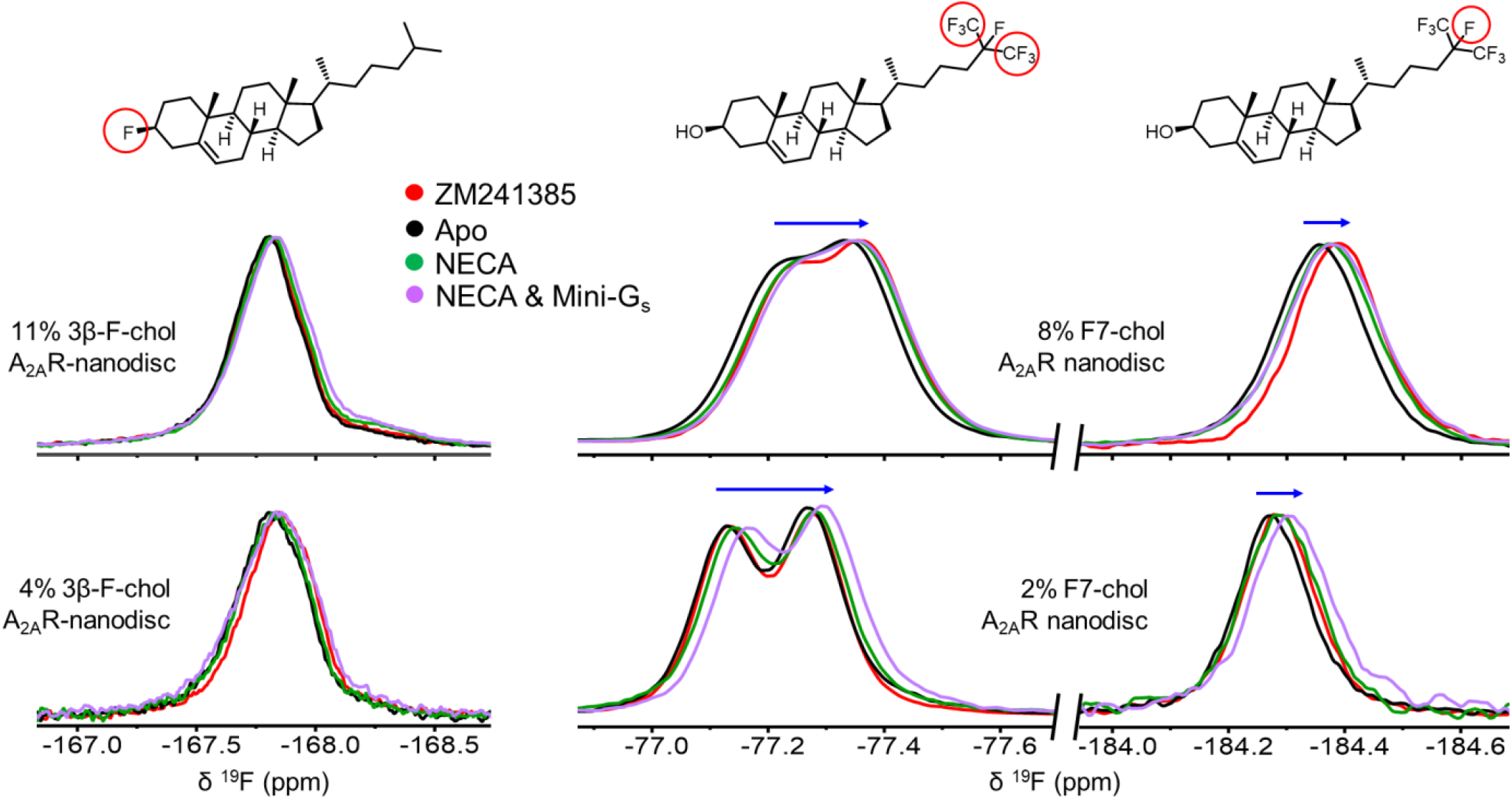
^19^F-cholesterol analogs behave similarly in the presence of agonist- and inverse agonist-bound A_2A_R. ^19^F NMR spectra of 3β-F-chol **(A)** and F7-chol **(B)** in A_2A_R-embedded nanodiscs in the presence of inverse agonist (ZM241385), no ligand, full agonist (NECA), or NECA + mini-G_s_. The fluorine groups giving rise to each resonance is shown above their corresponding peaks and the blue arrows indicate the direction of chemical shift change in response to the addition of ligand and miniG_s_.

## Materials and Methods

### A_2A_R expression, purification, and nanodisc reconstitution

Receptor cloning, expression, and purification have been described previously (Huang et al., 2021; Ye et al., 2016). Briefly, *Pichia pastoris* (*P. pastoris*) SMD 1163 (*Δhis4 Δpep4 Δprb1*) cells carrying the gene for A_2A_R (residues 2-317 with the V229C mutation for ^19^F-labeling) were grown to high density in either shaker flasks or a bioreactor. Methanol (5% v/v) was added every 12-16 h to induce expression and the cells were harvested after 60-72 h post induction. The receptors were extracted from the yeast membrane, reacted with the fluorine tag 2-Bromo-N-[4- (trifluoromethyl)phenyl] acetamide (BTFMA) when applicable, and further purified in the absence of cholesterol or cholesterol analogs. Prior to cholesterol incorporation, the receptors were reconstituted in rHDL nanodiscs using a 3:2 ratio of 1-palmitoyl-2-oleoyl-sn-glycero-3-phosphocholine (POPC) to 1-palmitoyl-2-oleoyl-sn-glycero-3-phospho-(1’-rac-glycerol) (POPG) and the MSPΔH5 membrane scaffold protein (Franz Hagn, Manuel Etzkorn, 2013). The sample was purified using a HiLoad 16/600 Superdex 200 preparatory grade size exclusion column equilibrated with nanodisc storage buffer (50 mM HEPES, pH 7.4, 100 mM NaCl), and the peak containing monodisperse nanodiscs were collected for cholesterol incorporation and further purification.

### Incorporation of cholesterol and cholesterol analogs

Incorporation of cholesterol and its fluorinated analogs in nanodiscs was achieved via incubation of the nanodiscs with cholesterol solubilized in methyl-β-cyclodextrin (MβCD, MilliporeSigma Canada, Oakville, Canada). One to two days prior to incorporation, a concentrated MβCD-cholesterol stock was prepared by mixing cholesterol (MilliporeSigma) with MβCD buffer (50 mM HEPES, pH 7.4, 100 mM NaCl, 40 mM MβCD) to a final concentration of 8 mM (4 mM in the case of fluorinated analogs, due to their increased hydrophobicity). The mixture was sonicated briefly to disperse any large chunks, then incubated at 30 °C for 24-36 hours with shaking until the solution is clear to the eye. The solution is filtered through a 0.2 µM filter to eliminate any undissolved particles, then diluted with MβCD buffer to make MβCD-cholesterol stocks containing 0.8 mM, 2 mM, 3 mM, and 4 mM cholesterol (0.8 mM and 3 mM in the case of fluorinated analogs). These stocks were mixed with nanodiscs collected from the size exclusion column described above (containing both empty and A_2A_R-embedded nanodiscs, which co-eluted) in a 1:3 v/v ratio, such that the final concentrations in the mixtures are 10 mM MβCD, 20-30 µM nanodisc, and 0.2 mM, 0.5 mM, 0.75 mM, or 1 mM cholesterol, for different levels of cholesterol incorporation. The mixtures were incubated at room temperature for 15 min with gentle shaking, then diluted 10-fold with nanodisc storage buffer containing 1-2 mL bed volume of Ni-NTA resin prior to incubation at 4 °C for 2 h. After incubation, Ni-NTA resins were collected using a gravity column and washed extensively with nanodisc storage buffer to remove residual empty nanodiscs, MβCD, and MβCD-cholesterol. Nanodiscs containing the His-tagged A_2A_R were eluted from the column using elution buffer (50 mM HEPES, pH 7.4, 100 mM NaCl, 250 mM imidazole), concentrated, and exchanged to nanodisc storage buffer for subsequent experiments. For empty nanodiscs, a His_6_-tagged MSPΔH5 protein was used. The reconstituted discs were treated with MβCD-cholesterol as above, incubated with Ni-NTA resins, and the MβCD was washed away prior to elution and concentration.

### Lipid quantification

Phospholipid concentrations were measured using a modified sulfo-phospho-vanillin assay (Frings and Dunn, 1970). Each sample containing unsaturated phospholipids (nanodiscs or phospholipid standards) was dissolved in 50-fold volume of concentrated sulfuric acid and incubated in a boiling water bath for 10 min. The samples were cooled in a cold-water bath for 5 min, then diluted 16-fold with a phospho-vanillin reagent (0.12% w/v vanillin dissolved in 68% v/v phosphoric acid). The samples were incubated in the dark for 30 min prior to absorbance measurements at 525 nm using a spectrophotometer. Lipid concentrations were determined using standard curves of A_525_ from pure POPC and POPG. In the case of nanodiscs, the lipid concentrations were determined using standard curves of both POPC and POPG:

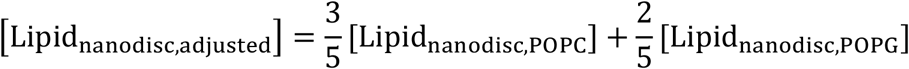

### Quantification of cholesterol and fluorinated cholesterol analogs

Cholesterol concentrations were measured calorimetrically using a commercial kit (R-Biopharm and Roche Diagnostics, Cat. No. 10139050035) following the manufacturer’s protocol. The concentrations of 3β-F-cholesterol and F7-cholesterol (Avanti Polar Lipids) were estimated via integration of ^19^F NMR resonances of the cholesterol analog in relation to a reference compound (fluoroacetate in the case of 3β-F-cholesterol and trifluoroacetate in the case of F7-cholesterol), where the relative signal loss in the reference peak due to shortened relaxation delay was corrected for. Percent cholesterol in a given sample was calculated as follows:

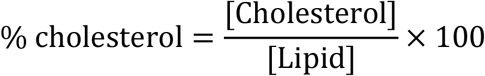

### G protein cloning, expression, and purification

The expression and purification of G_s_α, Gβγ, and mini-G_s_α have been described previously (Huang et al., 2021) with the only difference being that a wild-type G_s_α was used in the current work. To generate this construct, a double-stranded DNA fragment for the wild-type G_s_α short isoform was codon optimized and synthesized using the GeneArt service from Thermofisher. This fragment carried overlapping sequences with the previously described pET15b MBP-G_s_α mutant sequence (Huang et al., 2021). The plasmid was digested with XhoI and SacI (New England BioLabs, Ipswich, MA, USA) to remove the mutant G_s_α sequence and purified via electrophoresis and gel extraction kit (Bio Basic, Markham, Canada). The resulted plasmid backbone and DNA fragment were fused using the pEasy assembly kit from TransGen Biotech following manufacturer’s instructions. The plasmid was transformed into *Escherichia coli* (*E. coli*) BL21 (DE3) cells and a resulting colony containing the gene for the wild-type G_s_α was selected for protein expression.

### NMR experiments

NMR samples were prepared in nanodisc storage buffer with 20-100 µM A_2A_R-V229C (BTFMA-labelled for receptor NMR, unlabelled for ^19^F-cholesterol NMR), 10% D_2_O, and 20 µM sodium trifluoroacetate (TFA) or 100 µM fluoroacetate as the ^19^F chemical shift reference. For samples containing G protein (1.1-fold excess), the buffer also included 100 µM GDP, 2 mM MgCl_2_, and 5% glycerol. When applicable, A_2A_R ligands were added at saturating concentrations (1 mM NECA, 500 µM LUF5834, or 500 µM ZM241385). All samples were sterile-filtered and prepared in sterile Shigemi tubes to prevent microbial contamination. NMR experiments were acquired at 20 °C on a 500 MHz Varian Inova spectrometer equipped with a 5 mm room temperature inverse HFX probe. A typical fluorine NMR experiment included a 100 ms recycle delay, a 5.5 μs (45°) excitation pulse, and a 500 ms acquisition time. Each experiment acquired between 100,000-400,000 scans, yielding a S/N of approximately 50-100. Spectra were processed using MestReNova (Mestrelab Research S.L.) employing chemical shift referencing (−75.6 ppm for TFA and -217 ppm for fluoroacetate), baseline correction, zero filling, and exponential apodization equivalent to a 5-20 Hz line broadening. For high pressure NMR, the sample was transferred to a 3 mm zirconia tube (Daedalus Innovations, Aston, PA, USA) and covered with paraffin oil. The tube was placed inside a 600 MHz Varian Inova spectrometer equipped with a triple-resonance cryoprobe tunable to ^19^F, via a stainless-steel fluid line connected to an Xtreme-60 syringe pump (Daedalus Innovations) prefilled with paraffin oil as the pressurizing fluid. Pressure was increased at a rate of 100 bar/min to the desired value, and the sample was equilibrated for 5 min at the final set pressure prior to acquisition at 20 °C.

### Membrane fluidity measurements

A_2A_R-embedded nanodiscs were incubated with the fluorescent probe Laurdan (MilliporeSigma) at room temperature for 30 min in the dark at a final concentration of 1 μM A_2A_R and 10 μM Laurdan (diluted from a 10 mM dimethylformamide stock). Free Laurdan was removed by extensive buffer-exchange with the nanodisc storage buffer and subsequently filtering the sample through a 0.2 μm filter. Flow-through from the final round of buffer-exchange was kept for background correction. The samples were transferred to a black 384-well plate and the fluorescent emission spectra (410 nm – 520 nm) were acquired using a TECAN Spark multi-mode plate reader (Tecan, Männedorf, Switzerland) at 26 °C with an excitation wavelength of 350 nm. Each emission spectrum was background-corrected, then area-normalized to the 13% cholesterol condition. The generalized polarization (GP) of each sample was determined using the formula 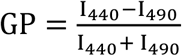, where I_440_ and I_490_ represent the emission intensities at 440 nm and 490 nm, respectively.

### GTP hydrolysis experiments

GTP hydrolysis experiments were carried out using the GTPase-Glo™ assay kit (Promega, Madison, WI, USA) following the manufacturer’s protocol (Mondal et al., 2015). Briefly, purified receptor and G protein were incubated at room temperature in a buffer containing 50 mM HEPES, pH 7.4, 100 mM NaCl, 2 mM MgCl_2_, 1 μM GDP, and 4 μM GTP, at a final concentration of 250 nM G protein, 250 nM A_2A_R, and various concentrations of the agonist NECA. Control reactions included buffer with GTP but in the absence of either A_2A_R or both A_2A_R and G protein. After 90 min, unreacted GTP was converted to ATP prior to the addition of a detection reagent containing luciferase. The resulting luminescence, which is proportional to the amount of unreacted GTP, was measured using a TECAN Spark multi-mode plate reader with an integration time of 1 min. GTP hydrolysis was determined as follows:

G protein only (in the absence of A_2A_R):

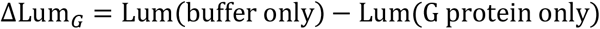

In the presence of A_2A_R:

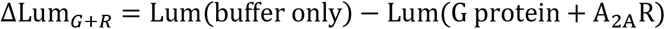

where Lum is the luminescence signal intensity.

The relative GTP hydrolysis for each A_2A_R (NECA) sample was calculated as follows:

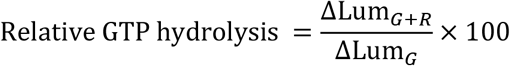

The NECA dose-response data were fit using a variable slope model in GraphPad Prism 8.4.2 employing the equation:

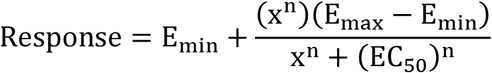

where x is the agonist concentration, E_min_ is the minimum response, E_max_ is the maximum response, EC_50_ is the agonist concentration that promotes half-maximum response, and n is the Hill coefficient.

### Dynamic Light Scattering

DLS samples were prepared in nanodisc storage buffer containing 5 μM A_2A_R-embedded nanodiscs supplemented with different mol% of cholesterol. Each sample was filtered through a 0.2 μm syringe filter to remove large dust particles before transferring to a small-volume 10 mm quartz cuvette (Starna Cells, Atascadera, CA, USA). DLS measurements were carried out inside a Zetasizer Nano-ZS particle size analyzer (Malvern Panalytical, Malvern, United Kindom) equipped with a He-Ne laser (λ = 633nm). Samples were equilibrated at 25 °C for 2 minutes and the scattered light was measured at a 173° backscatter angle. The resulting correlation function was analyzed using the general purpose (non-negative least squares) analysis model in the Zetasizer software (v7.13, Malvern Panalytical) for distribution analysis, assuming a buffer viscosity of 0.9066 cP, a buffer refractive index of 1.332, and a protein refractive index of 1.450. Data was averaged over three independent trials, each having 3 replicate measurements of 10-20 scans.

### Synthesis of 3β-fluoro-cholest-5-ene

3β-fluoro-cholest-5-ene was synthesized from cholesterol in one step, using the deoxyfluorination reagent DAST (diethylaminosulfur trifluoride, Toronto Research Chemicals, North York, Canada). Though fluorinations with DAST often proceed through an S_N_2 mechanism, fluorination of cholesterol is known to retain its stereochemistry (Rozen et al., 1979). This results from homoallylic participation forming a carbonium ion intermediate (Li et al., 2016; Rozen et al., 1979).

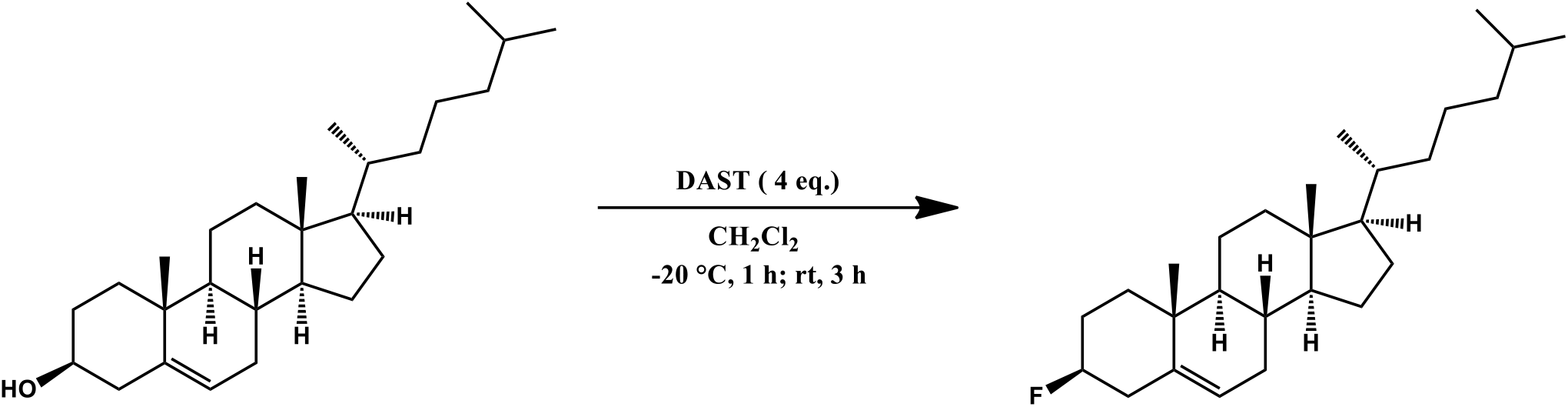

Cholesterol (650 mg, 1.68 mmol) was dissolved in dry CH_2_Cl_2_ (15 mL) in a plastic reaction vessel under argon. The mixture was cooled to -20 °C and DAST (4 eq., 0.89 mL) was added dropwise over 5 min. The solution was stirred at -20 °C for 1 h. The cooling bath was removed, and the reaction was continued at rt for 3 h. It was quenched by slowly pouring the mixture into a vigorously stirred solution of sodium bicarbonate at 0 °C. After the bubbling stopped, the aqueous phase was extracted twice with CH_2_Cl_2_ (50 mL). The organic layer was washed with brine and concentrated to give an orange syrup. Silica column chromatography (eluent: 100% pentanes, R_f_ = 0.27) yielded the product as a white solid (283 mg, 43%). The product’s spectroscopic characterization was consistent with published data (Li et al., 2016; Reibel et al., 2015).

^**1**^**H NMR** (400 MHz, CDCl_3_) δ 5.39 (d, *J* = 4.9, 1H), 4.67 – 4.13 (dm, ^2^*J*_H-F_ = 50.4 Hz, 1H), 2.44 (t, *J* = 7.0, 2H), 2.10 – 1.92 (m, 3H), 1.93 – 1.78 (m, 2H), 1.77 – 1.63 (m, 1H), 1.63 – 0.80 (m, 32H), 0.69 (s, 3H). ^**19**^**F NMR** (377 MHz, CDCl_3_) δ -167.82 (dm, ^2^*J*_F-H_ = 50.4 Hz). ^**13**^**C NMR** (101 MHz, CDCl_3_) δ 139.50 (d, *J* = 12.6 Hz), 123.16 (d, *J* = 1.3 Hz), 92.98 (d, *J* = 174.1 Hz), 56.88, 56.33, 50.17 (d, *J* = 1.8 Hz), 42.49, 39.91, 39.69, 39.57 (d, *J* = 19.3 Hz), 36.69 (d, *J* = 1.2 Hz), 36.53 (d, *J* = 10.8 Hz), 36.36, 35.95, 32.09 (d, *J* = 1.1 Hz), 32.03, 28.95 (d, *J* = 17.5 Hz), 28.39, 28.18, 24.45, 24.01, 22.98, 22.73, 21.29, 19.47, 18.89, 12.02. **HRMS (EI)**: Calcd. For C_27_H_45_F: 388.3505; Found: 388.3506.

### Computational rigidity-transmission allostery analysis

The fully active state of A_2A_R in complex with G_s_αβγ, NECA and GDP was constructed, equilibrated and relaxed in a 1 µs MD simulation in 4:1 POPC:cholesterol extended membrane as previously described (Huang et al., 2021). This model was used to probe agonist-induced allosteric communication in the A_2A_R-Gαβγ complex with rigidity-transmission allostery (RTA) algorithm, whose details have been previously described (Huang et al., 2021; Ye et al., 2018). The RTA algorithm is a computational method based on mathematical rigidity theory, which predicts how perturbations of conformational rigidity and flexibility (conformational degrees of freedom) at one site transmit across a protein or a protein complex to modify degrees of freedom at other distant sites (Sljoka, 2021). Here, RTA was applied to examine the allosteric pathways between the orthosteric pocket and distal regions in the A_2A_R-Gαβγ complex with and without cholesterol. We quantified the available conformational degrees of freedom at every residue before and after rigidification of the agonist NECA. The change in degrees of freedom was then extracted for each residue, which represents the extent of allosteric transmission from the orthosteric pocket. In the presence of cholesterols, the analysis was carried out as previously described (Huang et al., 2021). To measure the impact of cholesterol on allosteric communication, the same analysis was repeated upon removal of all seven cholesterols found within 6 Å of the receptor.

## Author contributions

S.K.H. and R.S.P. designed the research. S.K.H and O.A. performed protein expression and purification for A_2A_R. S.K.H. performed protein expression and purification for G_s_α and mini-G. A.P. and S.K.H conducted expression and purification of Gβγ. L.P cloned the wild-type G_s_α construct. S.K.H. and O.A. performed the NMR experiments. S.K.H. performed the GTP hydrolysis assays and DLS experiments. R.J.P. performed the Laurdan fluorescence experiments. Z.A.M. and M.N. provided the 3β-fluoro-cholest-5-ene. A.S. performed the RTA analysis. S.K.H and R.S.P prepared the manuscript. R.S.P. supervised the project.

## Acknowledgements

This work was supported by the CIHR Operating Grant MOP-43998 to R.S.P.; S.K.H. was supported by Alexander Graham Bell Canada Graduate Scholarship-Doctoral from NSERC; A.S. was supported by CREST, Japan Science and Technology Agency (JST), Japan JPMJCR1402; R.J.P. was supported by QEII F.E. Beamish Graduate Scholarship in Science and Technology. Special thanks to Dmitry Pichugin for NMR spectrometer maintenance.

